# ILC3s are Required for Enterocyte Homeostasis to Food Intake

**DOI:** 10.64898/2026.04.20.719606

**Authors:** Emelyne Lécuyer, Fabian Guendel, Sascha Cording, Giulia Nigro, Jasna Medvedovic, Sophie Dulauroy, Helena Moguel-Houssin, Marion Rincel, Benoit Chassaing, Francina Langa, François Déjardin, Gérard Eberl

## Abstract

Food provides nutrients that are selectively absorbed by the intestine, but, at the same time, may contain elements that challenge the intestinal barrier and induce post-prandial inflammation (PPI). How PPI is controlled in order to avoid pathological perturbation of homeostasis remains unclear. Here, we report that during fasting, enterocytes increase their absorptive potential and oxidative metabolism, a program that is largely reversed upon food intake of lipids that perturb the intestinal barrier and induce PPI. Such perturbation is countered by ILC3s, in the absence of which PPI increases, program reversal does not occur, and enterocytes engage into excessive oxidative metabolism. This enterocyte state leads to critical hypoglycemia as a consequence of decreased glucose absorption and increased insulinemia, recapitulating the pathological situation found in patients suffering from intestinal damage and sepsis. We hereby uncover a critical function for ILC3s in maintaining enterocyte homeostasis upon challenging food intake.

## INTRODUCTION

The function of the intestine is to process food and absorb nutrients for the organism. Absorption is performed by enterocytes within the single layer of intestinal epithelial cells (IECs), which form an essential component of the intestinal barrier. IEC secrete mucus, produce anti-bacterial peptides and translocate secretory IgA to buffer their interaction with the microbiota, regulate transcellular trafficking and translocate material from the lumen into the lamina propria for scrutiny by cells of the immune system (*1*).

Nevertheless, a lipid-rich diet destabilizes the mucus layers and increases intestinal permeability, leading to the passage from the lumen into the lamina propria and the bloodstream of microbial compounds, such as lipopolysaccharides (LPS), and the induction of systemic postprandial inflammation (PPI) (*2–5*). Such diet also favors the expansion of facultative aerobes, such as γ*- Proteobacteria*, which include species with a high inflammatory potential (*6*). Prolonged increase in PPI is a risk factor for chronic inflammatory pathologies, such as type 2 diabetes (*4*). However, the intestine faces such daily challenges generally without pathologic consequences (*7, 8*), but the mechanisms required to maintain intestinal and systemic homeostasis upon PPI remain largely unclear.

The intestine hosts the largest array of effector immune cells in the body, an activity that is induced by the colonization of the lumen by numerous and diverse types of microbes (*1, 9*). The bacterial and fungal microbiota, in particular, induce a robust type 3 immune response, carried by Th17 cells and type 3 innate lymphoid cells (ILC3) (*10*). These cells express the lymphotoxins LTα_1_β_2_ and LTα_3_, as well as IL-22, involved in the crosstalk with IECs and stromal cells to induce the production of tight junction proteins, anti-bacterial peptides, fucosylation of glycans protecting IECs (*11, 12*), and resistance of epithelial stem cells to apoptosis (*13, 14*). They also express IL-17 involved in the recruitment and activation of granulocytes (*15*). Consequently, the effector activity of Th17 cells and ILC3s increases resistance to bacterial and fungal infections, regulates the composition and confinement of the microbiota, and decreases endotoxemia (*16–18*). The activity of Th17 cells and ILC3s follows a circadian rhythm (*8, 19, 20*), associated with feeding and microbiota composition (*7*), the deregulation of which leads to elevated type 3 responses and susceptibility to inflammatory pathology (*19*). The effector functions of ILC3s are also potentiated by vasoactive intestinal peptide (VIP) induced by food intake (*21–23*).

Here, we report that ILC3s play a key and non-redundant role in the control of PPI and maintenance of IECs, intestinal and systemic homeostasis. While enterocytes increased their absorptive potential and oxidative metabolism during fasting, presumably to optimize energy production from limited food availability, they reverted this program during lipid-rich food intake while PPI and ILC3s activity reached their peak levels. In the absence of ILC3s, however, this program reversion did not occur and oxidative metabolism further increased, leading to oxidative stress in IECs, and consequent increase in intestinal permeability and PPI, defective glucose absorption and increased insulinemia, a pathologic metabolic condition that is found in the context of intestinal damage, infection and sepsis.

## RESULTS

### Fasting increases nutrient absorption and oxidative metabolism in IECs

In order to assess the impact of a lipid-rich food intake on the intestinal barrier and enterocyte function, and avoid variability in feeding behavior between individuals, mice were fasted overnight and then fed by gavage (fig. S1A). We thus first determined the transcriptional state of enterocytes at fasting in all segments of the small intestine (duodenum, jejunum and ileum) by single cell RNA sequencing. Eleven types of IECs were identified, in accordance with earlier reports (*24*), including four types of enterocytes (Fig. 1A). Fasting induced significant remodeling of the transcriptional landscape of enterocytes in all intestinal segments (Fig. 1B and S1B), reflected by a strong increase in the expression of genes involved in oxidative metabolism, such as fatty acid oxidation (FAO), oxidative phosphorylation (OxPhos) in the lower jejunum and ileum, in particular in genes encoded by mitochondrial DNA, and detoxification (Fig. 1C). Furthermore, a broad range of nutrient transporters were upregulated in all intestinal segments, reflecting an overall adaptation to low nutrient availability through an increase in transport and energy extraction through oxidative metabolism, as reported earlier at the level of intestinal stem cells (*25*). Interestingly, such metabolic adaptation to fasting was also observed, by quantitative RT-PCR of transcripts coding for nutrient transporters (*Scl40a1*, *Slc37a2*) and detoxifiers (*Cyp2c66*), during the fasting phase of the circadian rhythm (Fig. 1D) and in cultures of intestinal organoids in “fasting” low glucose conditions (fig. S1C).

**Fig. 1.**
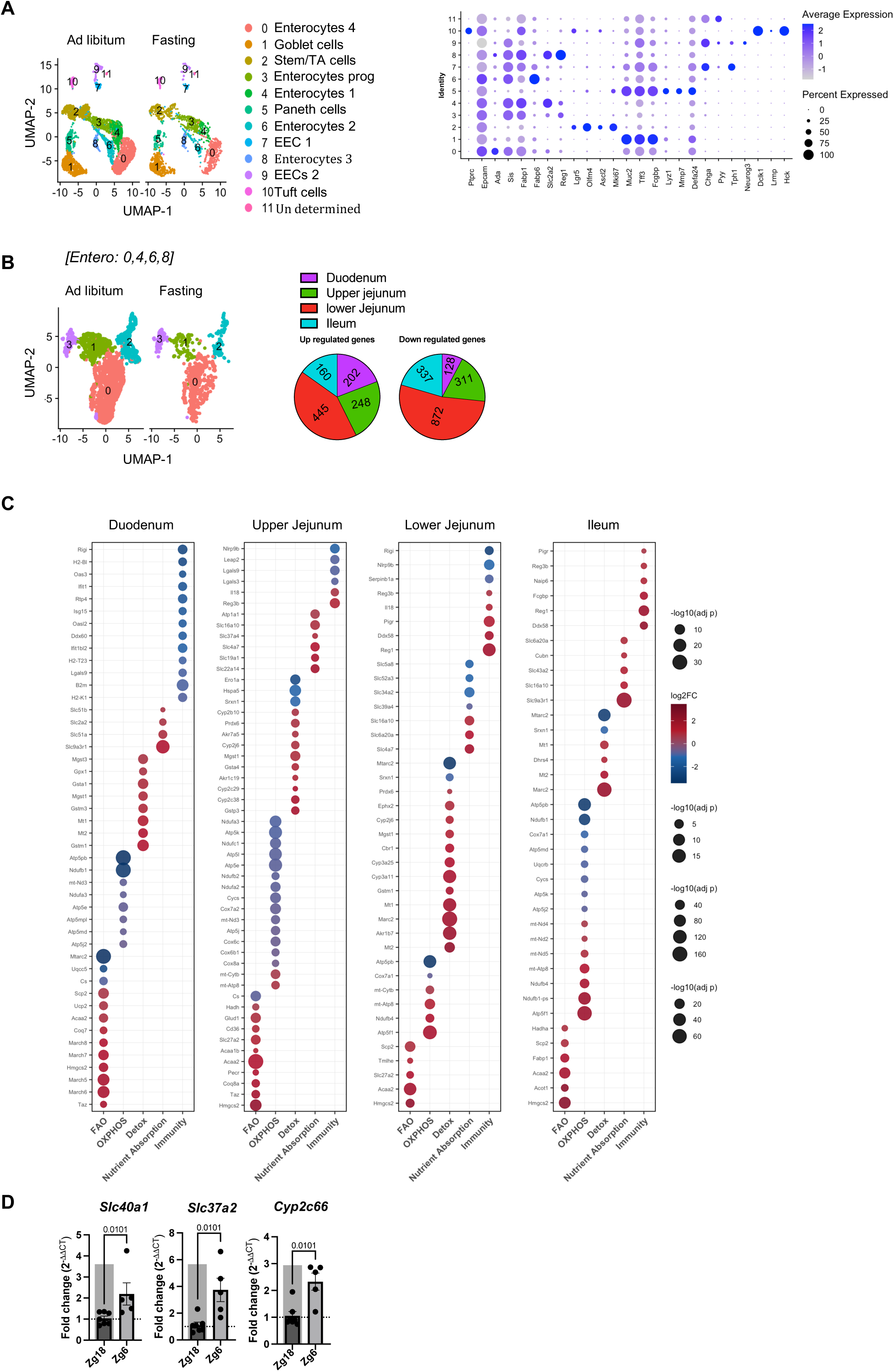
Fasting increases nutrient absorption and oxidative metabolism in IECs. **(A)** UMAP projection of scRNA-seq data from intestinal epithelial cells (IECs) isolated from control fasted mice (PBS) and control ad libitum-fed mice. Colors denote unsupervised clusters identified by Seurat; annotation was based on canonical intestinal epithelial cell markers (see Dot plot). (**B**) UMAP projection of enterocytes (clusters 0, 4, 6, and 8) from control fasted and ad libitum-fed mice, colored by Seurat-defined clusters. Clusters were annotated using canonical markers corresponding to distinct intestinal regions (fig. S1B). The accompanying pie chart depicts the proportions and numbers of up- and down-regulated genes identified by differential expression analysis comparing fasted and ad libitum conditions. (**C**) Dot plot showing representative up- and down-regulated DEGs between fasted and ad libitum groups. Genes were functionally categorized using an AI-assisted LLM-based classification across intestinal regions. (**D**) Relative expression of *Slc40a1*, *Cyp2c66*, and *Slc37a2* in IECs from ad libitum-fed mice (mean ± SEM; n = 5–6 mice per group) collected at 12 a.m. (Zeitgeber 18) and 12 p.m. (Zeitgeber 6). Data in (A-C) are representative of one single cell RNA sequencing experiment. Data in (D) represent a pool of three independent experiments and symbol indicates individual mice. *P* values were determined by two-tailed nonparametric Mann–Whitney U test. Exact *P* values are indicated on graphs.

### Lipid-rich food intake induces ILC3 activation and IEC decrease in oxidative metabolism

It was reported previously that lipid-rich diets and lipid-rich food intake perturb the intestinal barrier by altering the microbiota composition, IEC function and mucosal immunity (*2, 26–32*). To characterize more in details the impact of a lipid-rich food intake upon fasting, overnight fasted mice were fed coconut oil (enriched in saturated lipids), soybean oil (enriched in unsaturated lipids), linseed oil (enriched in unsaturated lipids), a glucose solution or a protein-rich bolus. Coconut oil significantly weakened the intestinal barrier (Fig. 2A), and induced maximal permeability 90 minutes after food intake (Fig. 2B), as well as an increase in γ*-Proteobacteria* (fig. S2A). The increase in gut permeability was associated with increased blood levels of IL-6 (Fig. 2C), a direct measurement of PPI (*4, 33*). The levels of IL-6 were dependent on microbiota and enterotoxins, as antibiotic treatment or the addition of γ*-Proteobacteria*-derived lipopolysaccharides (LPS) to the food intake decreased or increased such levels, respectively (Fig. 2D). Furthermore, the levels of intestinal permeability were correlated with the expression levels of *Il22* in the intestine (Fig. 2E), found to be produced exclusively by ILC3s (Fig. 2F and S2B), as well as elevated levels of *Il1b* (fig. S2C).

**Fig. 2.**
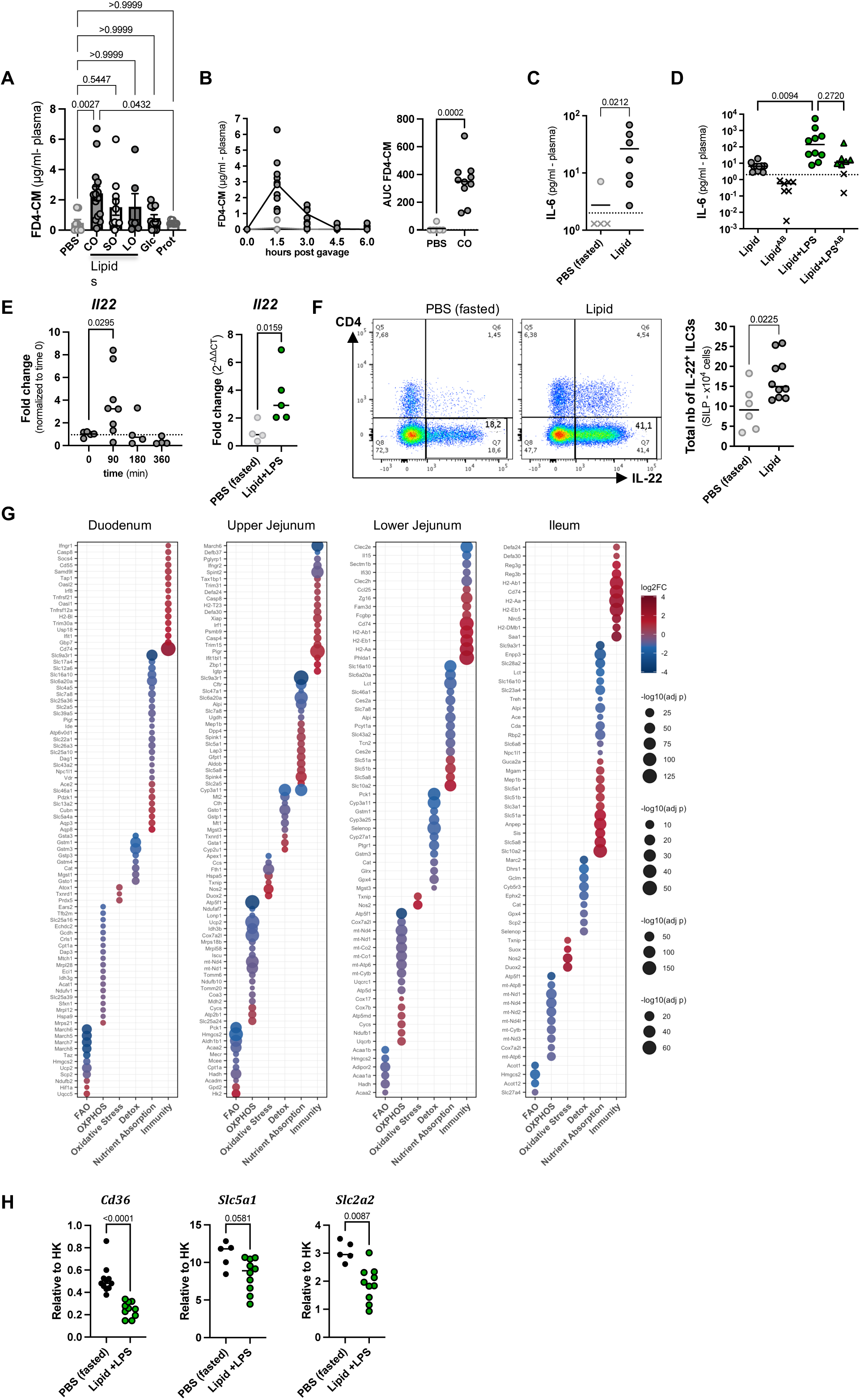
Lipid-rich food intake induces ILC3 activation and IEC oxidative adaptation. (**A**) Intestinal permeability (median; *n* = 6–15 per group) measured 90 min after intragastric gavage with PBS, coconut oil (CO), soybean oil (SO), linseed oil (LO), glucose (Glc), or protein (Prot). (**B**) *Left:* time course of intestinal permeability in PBS- (light gray) or CO-fed (dark gray) mice. *Right:* area under the curve (AUC) of the intestinal permeability test (median; *n* = 8–10 per group). (**C**) Plasma IL-6 concentration (median; n = 4–7 per group) in mice measured 90 min after CO (lipid) intake. Crosses (x) indicate mice below the linear detection limit of the assay. (**D**) Plasma IL-6 concentration (median; n = 7-12 per group) in mice fed CO (lipid) or CO (lipid)/LPS, with or without antibiotic (AB) treatment. Crosses (x) indicate mice below the linear detection limit of the assay. (**E**) *Left:* time course of *Il22* expression (median; *n* = 1–8 per group) in the ileum of mice following feeding with CO (lipid), as measured by qRT-PCR. Expression levels were normalized to the baseline (t0). The dotted line represents the mean expression level of PBS treated mice normalized to baseline. *Right: Il22* expression in ileal biopsies from mice 90 min after feeding with PBS (light gray) or CO (lipid)/LPS (green). (**F**) Representative dot plots of small intestinal ILC3s gated on viable CD45^low^RORgt^+^lin^-^ CD3^-^ CD127^+^ cells and quantification (median, n= 6 or 10 per group) of absolute numbers of IL-22^+^ ILC3s. (**G**) Dot plot of differentially expressed genes (DEGs) between CO (lipid)/LPS-fed and fasted (PBS) control mice. Genes were functionally categorized using an AI-assisted LLM-based classification across intestinal regions. (**H**) Relative expression of *Cd36*, *Slc5a1*, and *Slc2a2* measured by qRT-PCR in total IECs from control mice 90 min after dietary challenge with PBS (light gray) or CO lipid/LPS (green). Data in (A) to (F) and (H) are a pool of three to four independent experiments and representative of two or more independent experiments and symbol indicates individual mice. Data in (G) are representative of one single cell RNA sequencing experiment. *P* values were determined by two-tailed nonparametric Mann–Whitney U test or Kruskal–Wallis test. Exact *P* values are indicated on graphs.

In response to such lipid-rich food intake, enterocytes reversed their predominantly oxidative metabolism induced by fasting (Fig. 2G, fig. S2D). The expression of genes involved in FAO, OxPhos and detoxification was mostly downregulated, whereas the expression of genes involved in innate or adaptive immunity was increased, reflecting the increase in PPI. The expression of nutrients transporters was more nuanced, showing nonetheless a decrease in transcripts coding for lipid (*Cd36*) and glucose (*Slc5a1, Slc2a2*) transporters (Fig. 2H). The concurrent downregulation of metabolic and select transporter programs on the one hand, and the upregulation of immune transcripts on the other, suggests that enterocytes undergo a functional trade-off between nutrient absorption and barrier defense, likely required to maintain the integrity of the barrier upon food intake.

Together, these data show that a single lipid-rich food intake is sufficient to challenge the intestinal barrier, elicit PPI and reprogram IEC transcription from oxidative metabolism toward innate immunity. Furthermore, the kinetic association between IL-22 levels and barrier integrity, together with the exclusive expression of IL-22 by ILC3s, points to a central role for ILC3s in the IEC response to a feeding challenge.

### ILC3s are required for PPI control and enterocyte homeostasis

In order to assess the role of ILC3s during PPI, we developed a unique mouse model (RORγt-driven depletor of ILC3s, or ROD3) that allows for the rapid and specific ablation of ILC3s, thus preventing the immune system from developing compensatory mechanisms, as reported previously (*34*), in the short time frame of PPI. ROD3 mice express a floxed allele of the human diphtheria toxin (DT) receptor (DTR) under control of the *Rorc(ɣt)* promoter on a bacterial artificial chromosome (BAC) (*35*) (fig. S3A). This transgene allows for the ablation of ILC3 and ILC3-like cells (*10, 36–38*) without affecting T cells expressing the nuclear hormone receptor RORɣt, such as Th17 cells and bacteria- induced regulatory T cells (Tregs) (*39, 40*). When Rorc(ɣt)DTR (ROD) mice are crossed to Lck-Cre mice that express the Cre recombinase in T cells only (fig. S3B), the DTR sequence is excised in T cells, and only ILC3s express DTR (ROD3 mice). Of note, no ectopic expression of GFP reporting DTR was found in cells other than ILC3s or T cells (fig. S3C). As a consequence, treatment of ROD3 mice with diphtheria toxin (DT) led to the ablation of ILC3s but not of RORɣt^+^ T cells (fig. S3D). The ablation of ILC3s was nevertheless incomplete, possibly explained by the relative resistance of ILC3s to cellular stress, as reported upon irradiation or chemotherapy (*13, 14, 41*), or by ectodomain shedding of the DTR by TACE (*42*). However, surviving ILC3s showed reduced production of IL- 22, in line with the inhibition of protein translation by DT (fig. S3E).

As expected, expression of *Il22* in response to the dietary lipid-rich food intake was markedly reduced in the absence of ILC3s in DT-treated ROD3 mice (Fig. 3A). Loss of ILC3 function was also associated with increased gut permeability, elevated levels *Il6* and *Tnf* expression in the intestine (Fig. 3B), but not in DT-treated DEREG mice that deplete regulatory T cells (*43*), as a control of DT-induced cell death-mediated inflammation (fig. S3F). In this context, the expression of genes involved in FAO was not reduced but increased, as compared to mice with a normal complement of ILC3s, as was the expression of genes involved in oxidative stress, in particular in the jejunum (Fig. 3C). The increased expression of transcripts coding for the reactive oxygen radicals (ROS)-producing enzymes *Nox1*, *Nos2* and *Sod1* was confirmed by qRT-PCR (Fig. 3D), an increase found to be independent of T cells (fig. S3G), thus confirming the non-redundant role of ILC3s in the regulation of the oxidative stress response in IECs. Furthermore, blockade of IL-22 with anti-IL-22 neutralizing antibody following the lipid-rich food intake led to an accumulation of ROS species in IECs (Fig. 3E), confirming the key role of ILC3-derived IL-22 in this regulation.

**Fig. 3.**
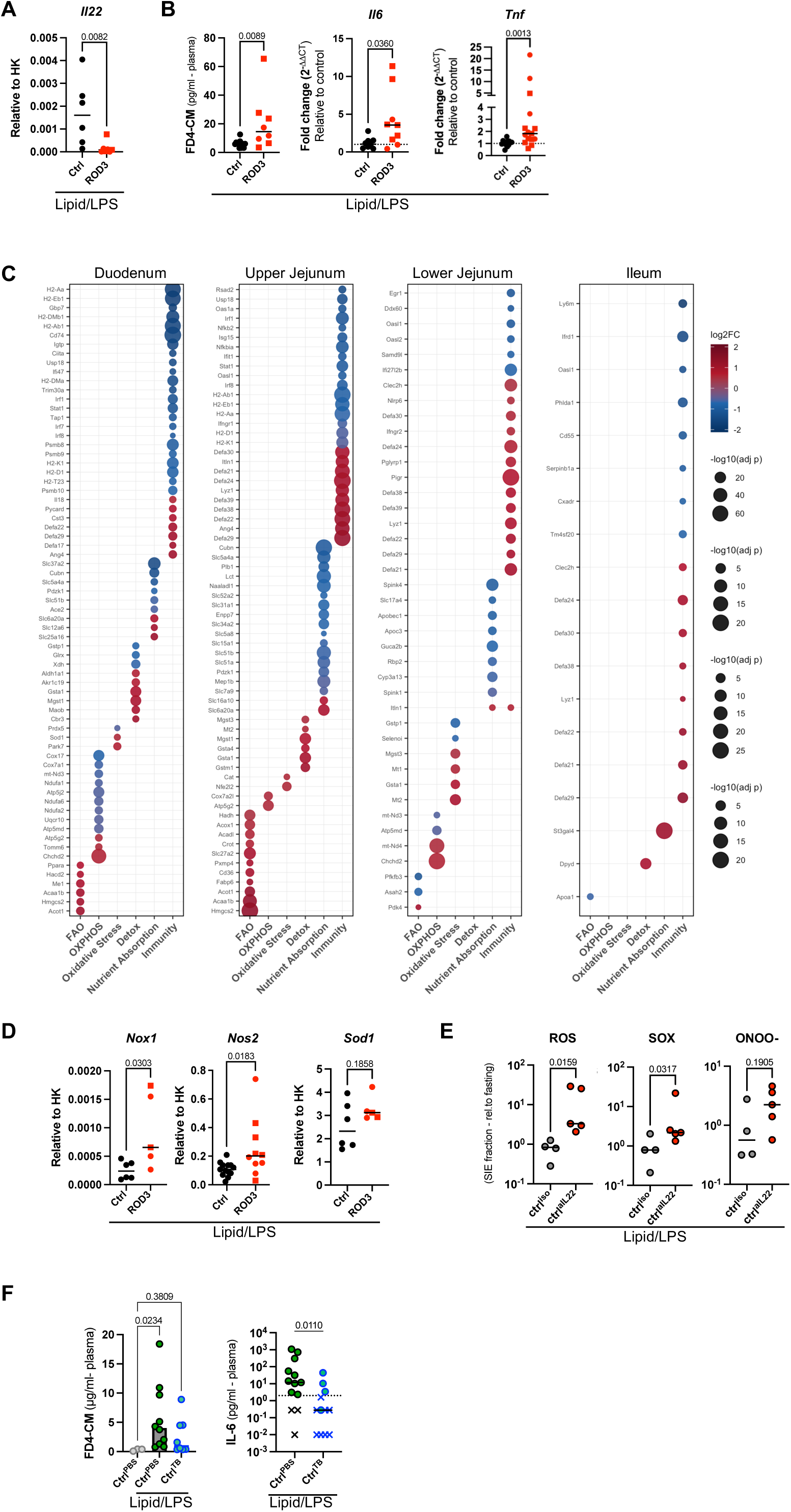
ILC3s are required for PPI control and enterocyte homeostasis. (**A**) Relative expression of *Il22* (median; n = 6–7 per group) determined by qRT-PCR on ileal biopsies from DT-treated control and ROD3 mice 90 min after feeding with CO lipid/LPS. (**B**) *Left:* Intestinal permeability measured by plasma FD4-CM levels (median; n = 8–12 per group), middle: *Il6* expression (median; n = 8–9 per group) and, *right: Tnf* expression determined by qRT-PCR in small intestinal epithelial cells from DT-treated control and ROD3 mice 90 min after feeding with CO lipid/LPS. (**C**) Dot plot of differentially expressed genes (DEGs) between DT-treated CO lipid/LPS-fed ROD3 and control mice. Genes were functionally categorized using AI-assisted LLM-based classification across intestinal regions. (**D**) Relative expression of *Nox1*, *Nos2*, and *Sod1* (median; n = 5–13 per group) determined by qRT-PCR in small intestinal epithelial cells from DT-treated control and ROD3 mice 90 min after feeding with CO lipid/LPS. (**E**) ROS, SOX and ONOO- levels on IEC 90 min after lipid/LPS feeding of mice that were pre-treated with anti-IL-22 neutralizing antibody or isotype control. (**F**) Intestinal permeability (median; n = 3–10 per group), and plasma IL-6 concentration (median; n = 3–10 per group), in PBS- (light gray), CO lipid/LPS-fed (dark gray-green), or CO lipid/LPS-fed mice pre-treated with tributyrin (light blue) 90 min after feeding, with or without tributyrin pre-treatment (administered 30 min before challenge). Data in (A) to (B) and (E) to (F) are a pool of two (E) to three independent experiments representative of one (E), two or more independent experiments, and symbol indicates individual mice. Data in (C) are representative of one single cell RNA sequencing experiment, in which two mice per group (aged 8 and 13 weeks, both sexes) were pooled together. *P* values were determined by two-tailed nonparametric Mann–Whitney U test or Kruskal–Wallis test. Exact *P* values are indicated on graphs.

The impact of oxidative stress on the intestinal barrier was demonstrated by the treatment of mice with tributyrin (TB), a triglyceride metabolized into butyrate and glycerol by host and microbial lipases (*44, 45*). As expected from previous reports (*46*), TB reduced gut permeability and PPI (Fig. 3F). These data show that, in the absence of ILC3s, the level of PPI is increased and enterocytes fail to revert their elevated oxidative metabolism, induced by fasting, to normal levels, upon lipid-rich food intake.

### PPI in the absence of ILC3s leads to defective glucose absorption and hypoglycemia

Oxidative stress in enterocytes affects the absorption and processing of dietary lipids (*47*) and carbohydrates (*48*). Accordingly, the increased levels of oxidative stress induced by lipid-rich food intake in the absence of ILC3s (Fig. 3C, D) led to lower levels of glycemia (Fig. 4A) in the absence of increased insulinemia (Fig. 4B). We assessed the glucose absorption potential of enterocytes and found lower expression of *Slc5a1* and *Slc2a2*, coding for the apical glucose transporter SGLT-1 and the basolateral glucose transporter GLUT2, respectively (Fig. 4C). The expression of *Slc5a1* and *Slc2a2* was strongly increased by fasting in the duodenum, and most affected in that segment by the absence of ILC3s (Fig. 4D). As a consequence, the duodenum did not absorb the traceable glucose analogue 2-NBDG in the absence of ILC3s (Fig. 4E) or of its effector cytokine IL-22 (Fig. 4F), whereas no such effect was observed in T cell-deficient mice (fig. S4A, B). Inhibition of oxidative stress with TB (Fig. 4G) in the absence of ILC3s or IL-22 effectively restored 2-NBDG absorption (Fig. 4E, F).

**Fig. 4.**
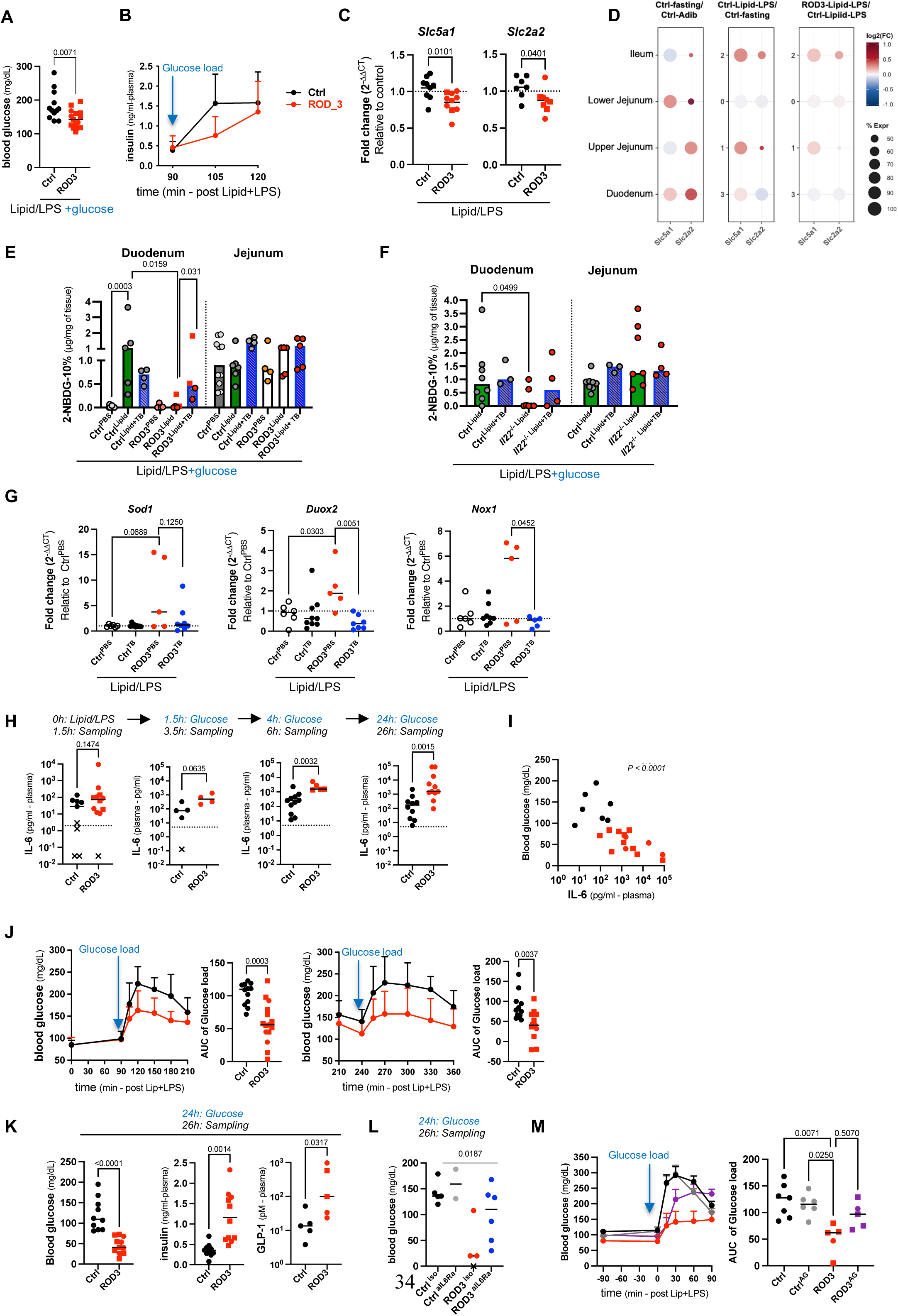
PPI in the absence of ILC3s leads to defective glucose absorption and hypoglycemia. (**A**) Blood glucose levels (median, n =12-15 per group) in control mice and ROD3 mice 15 mins after oral gavage of glucose solution following CO lipid/LPS feeding. Circles indicate ROD3 mice, and squares indicate ROD mice. (**B**) Plasma insulin levels (Mean ± SD, n = 4-8 per group) in control and ROD3 mice at the indicated time after oral gavage of glucose solution following CO lipid/LPS feeding. (**C**) Relative expression (median, n = 7-10 per group) of *Slc5a1*and *Slc2a2* as determined by qRT-PCR in intestinal epithelial cells from control and ROD3 mice 90 min after feeding with CO lipid/LPS. Circles indicate ROD3 mice, and squares indicate ROD mice. (**D**) Dot plot of fold changes in *Slc5a1* and *Slc2a2* expression in the following comparisons: control fasted versus ad libitum-fed mice; control mice fed CO lipid/LPS versus control fasted mice; and ROD3 mice fed CO lipid/LPS versus control mice fed CO lipid/LPS. (**E**) Absorption (median, n = 4-10 per group) of 2-NBDG (2- (*N*-(7-nitrobenz-2-oxa-1,3-diazol-4-yl)amino)-2-deoxyglucose) in the indicated intestinal compartment 15 min after administration in control and ROD3 mice previously fed and treated with PBS, CO lipid/LPS, PBS+Tributyrin (TB) or CO lipid/LPS+Tributyrin. Circles indicate ROD3 mice, and squares indicate ROD mice. (**F**) Absorption (median, n = 4-10 per group) of 2-NBDG in the indicated intestinal compartment 15 min after oral gavage in control mice and *Il22*^-/-^ mice previously fed and treated with PBS, CO lipid/LPS, PBS+Tributyrin (TB) or CO lipid/LPS+Tributyrin. (**G**) Relative expression (median, n= 5-9 per group) of *Sod1, Duox2*, *Nox1* as determined by qRT-PCR in small intestinal epithelial cells from DT-treated control mice and ROD3 mice 90 min after CO lipid/LPS feeding, or pre-treated with TB 30 min prior to the feeding. (**H**) Concentration of plasma IL-6 in control and ROD3 mice at indicated times after oral glucose administration. (**I**) Correlation between blood glucose and plasma IL-6 concentration in DT-treated control and ROD3 mice, 2 hours after oral glucose administration performed 24h after CO lipid/LPS feeding. (**J**) Blood glucose levels over time (Mean ± SD, n = 13-15 per group) and AUC (median n = 13-15 per group) in DT-treated control and ROD3 mice, 90min (left) and 4h (right) after CO lipid/LPS feeding. Circles indicate ROD3 mice, and squares indicate ROD mice. (**K**) Glycemia (left), insulinemia (middle) and total plasma GLP-1 levels (right), 24h after the CO lipid/LPS feeding. (**L**) Blood glucose levels in control and ROD3 mice treated with isotype control (iso) or anti-IL6Ra antibody (aIL6Ra) 2 hrs after glucose challenge performed 24h after CO lipid/LPS feeding. (**M**) Left: blood glucose levels over time (Mean ± SD, n = 5-7 per group) and, right: AUC (median, n = 5-7 per grouop) in DT-treated control and ROD3 mice, pre-treated with aminoguanidine (AG) for 16h and 90min after CO lipid/LPS feeding. Data in (A) to (C) and (E) to (M) are a pool of two to three experiments and representative of two or more independent experiments, and symbol indicates individual mice. Data in (D) are representative of one single cell RNA sequencing experiment, in which two mice per group (aged 8 and 13 weeks, both sexes) were pooled together. *P* values were determined by two-tailed nonparametric Mann–Whitney U test or Kruskal–Wallis test or Sperman correlation (I). Exact *P* values are indicated on graphs.

We next assessed whether oral administration of glucose restored normal glycemia, despite defective absorption by the duodenum. However, glucose administration further increased oxidative stress, PPI and the expansion of γ*-Proteobacteria* (Fig. 4H, S4C), and decreased glycemia (Fig. 4I, J), while the levels of insulin and GLP-1 increased (Fig. 4K), in accordance with a previous report showing that oral glucose promotes PPI and stimulates insulin production (*49*). Along these lines, normal levels of glycemia were restored by the inhibition of IL-6 binding to its receptor IL-6Ra (Fig. 4L). The treatment of mice with the NOS2 inhibitor aminoguanidine also restored normal glycemia (Fig. 4M), in accordance with the reported effect of glucose on enterocyte oxidative stress (*50*), altogether suggesting the establishment of a pathological causal loop between oxidative stress, increased PPI, glucose absorption, hypoglycemia and insulinemia.

Collectively, our data show that ILC3s are required for reducing oxidative metabolism in enterocytes after a PPI-inducing lipid-rich food intake. In the absence of ILC3s, increased oxidative stress in enterocytes increases gut permeability and PPI, and inhibits glucose absorption, together leading to a pathogenic loss in systemic glucose homeostasis. Thus, ILC3s play a key role in the homeostasis of enterocytes in the face of variable and potentially damaging dietary conditions.

## DISCUSSION

Th17 cells and ILC3s have been increasingly recognized as key players in modulating the intestinal adaptation to concurrent challenges in nutrient absorption and microbial pathogens (*21, 22, 51, 52*). These challenges follow a circadian rhythm (*7, 22*) and induce a transient postprandial inflammation (PPI) characterized by increased levels of serum IL-6 (*4*). Such challenges can occasionally become more acute depending on the lipid and carbohydrate composition of the food (*33*), and the presence of immunogenic bacteria in the intestine (*53*). Our study unveils a critical role for ILC3s in the regulation of intestinal epithelial adaptation to food intake, by preventing oxidative stress in enterocytes, maintaining physiological absorption of glucose, as well as maintaining an efficient intestinal barrier.

ILC3s play a critical role in intestinal homeostasis at multiple levels. A subset of ILC3s, the lymphoid tissue-inducer (LTi) cells, induce the development of Peyer’s patches in the late embryo and of isolated lymphoid follicles in the newborn (*54*). After birth, ILC3s contribute to intestinal anti- microbial and homeostatic responses by the production of IL-22 and, thereby, the expression of anti- bacterial peptides by IECs (*55*) and the amplification of anti-viral responses (*56*). ILC3s also produce lymphotoxin that induces protective fucosylation of IECs (*11*) and the activation of dendritic cells (*57, 58*). These functions are shared with Th17/22 cells, even though the TCR-independent activation of ILC3s allows them to respond promptly to inducer cytokines, such as IL-1β and IL-23, the former being induced by food intake and involved in the regulation of postprandial glycemia (*49*). Food intake also induces the expression of the vaso-intestinal peptide (VIP), which in turn potentiates the effector functions of ILC3s (*21–23*). Our data now show that the activation by food intake of ILC3s and their production of IL-22 are key to maintain intestinal homeostasis during a dietary challenge.

To more precisely assess the role of ILC3s in the control of PPI, we have developed a new mouse model for the specific and inducible ablation of ILC3s. Numerous mouse models have been developed to assess the functions of ILCs and ILC subsets, mostly based on the cell-specific deletion of key developmental factors (*59, 60*). However, a caveat of such cell specific-deficient mice is adaptation of the immune system to the loss of that cell subset. We have reported that the absence of ILC3s and Th17 cells in RORγt-deficient mice was indeed complemented in the intestine by the increased production of IgM and IgG (*34*). The timed ablation of ILC3s in ROD3 mice thus allows assessing the specific role of ILC3s during short experimental time windows that do not allow for complementation by other elements of the immune system. Using this system, we show here that the acute loss of ILC3s leads to a failure to maintain IEC homeostasis upon a challenging food intake.

We find that IECs increase their nutrient absorption potential and oxidative metabolism during fasting, presumably to maximalize the extraction of constituents and energy while nutrients are scarce (*61*). As expected, this adaption of IECs to fasting is reversed during (lipid-rich) food intake. We show that this program reversal is not accomplished in the absence of ILC3s, leading to excessive oxidative metabolism and oxidative stress in IECs. Different mechanisms could be involved in this failure to maintain IEC homeostasis in the absence of ILC3s. First, ILC3s, mostly through their production of IL-22, contribute to the control of the intestinal microbiota (*20, 62–64*). In the absence of ILC3s, we observe an increase in γ*-Proteobacteria* that in turn may contribute to the increased PPI through the production of highly immunogenic components, such as LPS, and increase IEC oxidative stress (*45, 46*). Second, the high level of IL-22 produced by ILC3s after lipid-rich food intake, which is coincident with the peak intestinal permeability (90 min after food intake), engages an intracellular signaling pathways that leads to the activation of STAT3. Interestingly, STAT3 stabilizes the mitochondrial electron transport chain and reduces ROS production (*65–67*). Such direct impact of ILC3 function on the regulation of oxidative stress may be further fueled by their LKB1-mediated sensitivity to such stress (*68*), in the absence of which IEC homeostasis may be further impaired. Interestingly, it has been recently reported that prolonged exposure to high fat diet eventually leads to a loss in ILC3s as a consequence of intracellular lipid accumulation and mitochondrial stress (*69*).

We report that increased oxidative stress impairs glucose absorption by IEC, which is restored by the treatment of mice with tributyrin (*45*). Oxidative stress decreases the expression of *Slc5a1* and *Slc2a2*, coding for the apical glucose transporter SGLT-1 and the basolateral glucose transporter Glut2, respectively. It was previously shown in cultures of Caco-2 cells that oxidative stress indeed limits glucose absorption (*48*), although the mechanisms involved remain unclear. Furthermore, we show that oral glucose supplementation further decreases glycemia, and increases PPI, in accordance with a previous reports showing that oral glucose promotes PPI, decreases glycemia and stimulates insulin production (*49*). From experiments on Caco-2 cells, it was proposed that glucose increases oxidative stress through glycation of cytoplasmic proteins (*50*). In addition, in the context of increased PPI and oxidative stress in IECs, it is possible that oral glucose further promotes oxidative stress by promoting anaerobic glycolysis and rapid ATP production to control for the expansion of pro- inflammatory γ*-Proteobacteria*, as discussed earlier (*45, 46*).

Altogether, a lipid-rich food intake in the context of ILC3 deficiency leads to positive feedback on IEC oxidative stress, intestinal permeability and inflammation, and, as a consequence, to a dangerous loss in glycemic homeostasis. Such conditions may be met during intestinal infection, inflammation and sepsis, during which the homeostatic function of ILC3s and associated cells may be overwhelmed (*69–71*), leading to a loss in IECs, gut and glycemic homeostasis. From our observations, and as proposed earlier (*45*), administration of compounds that regulate IEC oxidative stress may be key to prevent potentially lethal outcomes.

## MATERIALS AND METHODS

### Mice

Males and females 10- to 12-week-old specific pathogen free (SPF) C57Bl/6 were purchased from Charles River company. Males and females 7- to 16-week-old ROD^+/+^, ROD^Tg^/^+^, ROD^Tg^/^Tg^ (ROD), Lck-Cre^Tg/+^, ROD^Tg/Tg^x LckCre^Tg/+^ (ROD3) mice used for the experiments were bred and maintained at the Pasteur Institute breeding animal facility. In all experiments, males and females littermates have been used, and diphtheria toxin (DT)-injected ROD double negative (ROD^+/+^), DT-injected LckCre^Tg/+^, DT-injected ROD single positive transgenic (ROD^Tg/+^) or DT-injected ROD3 single positive transgenic (ROD^Tg/+^ x LckCre^Tg/+^) littermates were used as controls. RORγt-deficient mice (Rorc(gt)-/-), Rag1-Cre, Rosa26-YFP, DEREG, *Cd3*^-/-^ deficient and *Il22*^-/-^ mice were bred and maintained at the Pasteur Institute breeding animal facility. All the mice were housed at the Pasteur Institute animal facilities in a specific pathogen-free (SPF) environment, maintained under a 12-hours light-dark cycle (7 a.m. to 7 p.m.) unless otherwise indicated and given a standard chow diet (Safe) and neutral water (pH 6.5 to 7.5) ad libitum unless otherwise indicated. Experimental groups were both age- and sex-matched. Animal experiments were approved by the committee on animal experimentation of the Pasteur Institute and by the French Ministry for Higher Education, Research and Innovation.

### Mouse treatments

To examine the postprandial (PP) response to dietary challenges, 16h-fasted mice were fed intragastrically with a solution of pre-warmed (37°C, to allow melting) coconut oil (Sigma), soybean oil (Sigma), glucose (CDM Lavoisier), dietary protein solution (Sigma) or saline solution (5µl/g of body weight). When mentioned, the administration of coconut oil (CO) was immediately followed by an intragastric administration of LPS (10mg/kg) (Sigma). The antibiotic treatment consisted in one single dose of streptomycin (Sigma) (20mg per mouse) administered intragastrically, followed by a ciprofloxacin treatment (Sigma) administered in the drinking water (4mg/L) for one week prior to the dietary challenge. Diphtheria toxin (DT) (Sigma) was given intraperitoneally (50ng/g of body weight), following the timeline shown one day before the analysis of the mice. Blood glucose levels were determined with a glucose meter (Accu Check PERFORMA, Roche/ FreeStyle Optium Neo, Abbott) on blood samples collected from the tip of the tail vein. Larger volume of blood was collected by cardiac puncture, placed in EDTA pre-coated tubes (Sarstedt) for plasma analysis of circulating markers. Tributyrin (Sigma) was administered intragastrically (200µl per mouse) 30 min prior PP feeding. For IL-6R blocking experiments, mice received three intraperitoneal injections of antibody targeting the IL-6Rα chain (BioLegend), or isotype control (1mg/mouse in total: day-1PM:250µg, day0AM:250µg, day0PM:500µg). For IL-22 blocking experiments, mice received an intraperitoneal injection of 150µg of IL22-JOP (eBioscience) or isotype control at the initiation of fasting. Aminoguanidine (Sigma) was administered in the drinking water (0.5g/L) when starting the fasting period of the dietary challenge.

### Biochemical analysis of blood markers

For total GLP-1 measurement, blood samples were harvested in DPP4 inhibitor (10µl/ml of blood) (Sigma), placed in pre-coated EDTA tubes (Sarstedt) for plasma collection. Circulating concentrations of IL-6 and total GLP-1 were determined by Enzyme-Like Immunosorbent Assay (ELISA) kits (eBioscience, Sigma).

### Bacterial DNA extraction and quantitative real-time PCR

Cecal content or feces were harvested and snap frozen in liquid nitrogen and stored at −80°C. Bacterial DNA extraction was performed using the QIAamp PowerFecal Pro DNA Kits (Qiagen) according to manufacturer’s instructions. The primer pairs used in this study are listed:

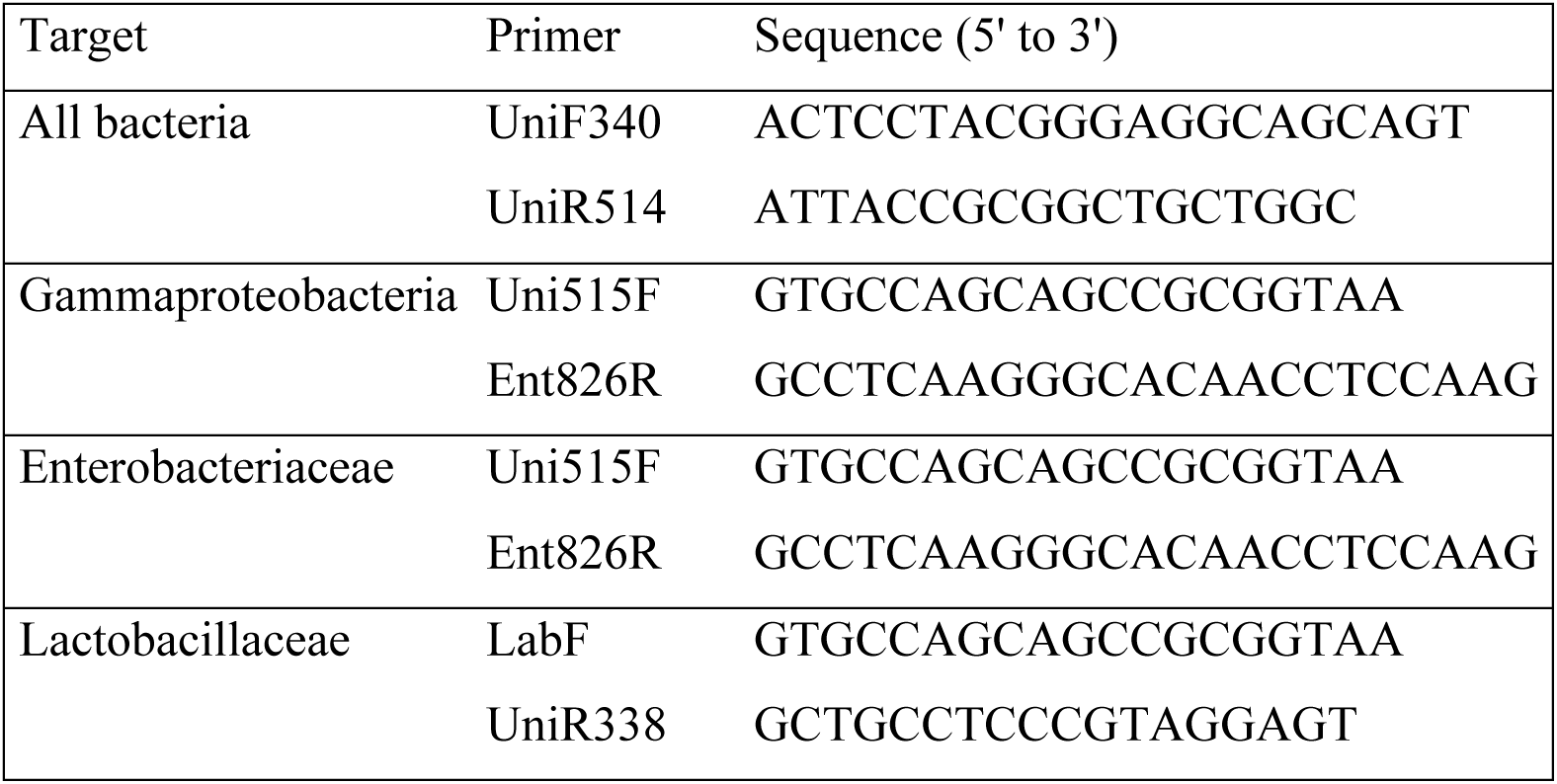

### Cell preparation for small intestinal lamina propria immune cells

After surgical removal of the small intestine and flushing of the luminal content with PBS and Peyer’s patches excision with scissors, the single cell suspension was obtained using the Lamina Propria Dissociation Kit (Miltenyi) following the manufacturer’s instructions. Leukocytes enrichment was then performed through a 40/80% (w/v) Percoll density gradient (GE Healthcare) centrifuged for 15min at 1900g at RT. Single cell preparations were washed prior further utilization.

### Cell preparation for small intestinal epithelial fraction

After surgical removal of the small intestine (SI) and flushing of the luminal content with PBS and Peyer’s patches excision with scissors, SI was longitudinally opened and cut into small pieces (5mm). SI pieces were placed in 2 baths of HBSS with EDTA (5mM) on ice with shaking every 20min and cell filtering on cell strainer 100µm. Supernatants were removed by centrifugation at each step (400g, 4°C, 10min). Epithelial cell fraction was then resuspended in Trizol (Invitrogen), snap frozen and kept at −80°C for further analysis.

### Cell stimulation before intracellular staining

Single cell suspensions were prepared from small intestinal lamina propria and resuspended in complete media containing DMEM-Glutamax, 8% fetal calf serum (FCS), HEPES (10 mM), 2- mercaptoethanol (0.05 mM), penicillin and streptomycin (100 U/ml) (all from Gibco). For ILC3 detection, cells were stimulated with IL-23 (40ng/ml; R&D Systems) and IL-7 (10ng/ml; R&D Systems) for 3h, and brefeldin A (10µg/ml; eBioscience) was added during the last 1.5h of culture. For Th17 and Th22 characterization, cells were stimulated for 3h with Phorbol 12-Myristate 13- Acetate (50 ng/ml; Thermo Scientific Chemicals) and ionomycin (500ng/ml; Invitrogen), adding brefeldin A (10µg/ml; eBioscience) for the last 1.5h of culture.

### Immunostaining and FACS analysis

Cells were preincubated with Fc-Block and LIVE/DEAD fixable blue dead stain kit (Invitrogen) was used prior to surface staining. Cells were surface stained for 20 min with the following antibodies: CD19-BV650/PE/PE-Cy7 (eBIO1D3), TCRb-PE-Cy7 (H57-597), CD11b-PE-Cy7 (M1/70), F4/80- PE-Cy7 (BM8), TER119-PE-Cy7 (TER-119), CD117-APC-AF760 (2B8), NKp46-biotin (29A1.4), CCR6-PE (140706), CD3ε-FITC (145-2C11), CD3ε-eFluor450/-APC-e780/- PECF394 (17A2; BioLegend), CD4-eFluor450 (GK1.5) or CD4-BV605/-APC (RM4-5; BD), CD8a-PE-Cy7 (53-6.7), Thy1.2-PE-Cy7/BV605/PerCP-eFluor710 (53-6.7), CD127-BUV737 (PC61.5), CD45.2-V500 (104; BD), CD45R-APCe780 (RA3-6B2), and streptavidin-PerCP-Cy-5.5 (eBioscience) or -AF647 (Invitrogen) were used for the detection of biotinylated antibodies. For intracellular staining, cells were fixed and permeabilized with a commercially available fixation/permeabilization buffer (eBioscience). Intracellular staining was performed with the following antibodies: RORγt-PE or RORγt-APC (AFKJS-9), IL-17A-PE or IL-17A-APC (TC11-18H10; BD), IL-22-PE (1H8PWSR), and polyclonal anti-GFP/YFP (A11122, Invitrogen). All antibodies were from eBioscience unless stated otherwise. Cytometry data was performed on a FACS Fortessa X20 and FACS Fortessa (BD Bioscience) with a Diva software. Data analysis of flow cytometry data was conducted with FlowJo Software V10.10.

### Intestinal permeability assay

Intestinal permeability measurement in vivo was based on the intestinal permeability to 4 000 Da Fluorescein isothiocyanate–Carboxymethyl–Dextran (FD4-CM) (Sigma-Aldrich). FD4-CM was administered intragastrically to mice (600 mg per kg of body weight). Post-gavage, blood was collected from the tip of the tail vein (40µl) into EDTA-coated tubes and centrifuged (4°C, 2000 g for 10 min). Plasma was diluted 1:10 (v/v) in phosphate buffered saline (PBS, pH 7.4) and the FD4- CM concentration was determined using a fluorescence plate reader (SLT, BMG Labtech) at 485 nm excitation and 535 nm emission wavelengths. Standard curves were obtained by diluting FD4-CM in non-treated plasma prepared in PBS (1:10 v/v).

### Oral glucose administration

1.5 g glucose per kg of body weight of a glucose solution (CDM Lavoisier) dissolved in sterile saline was administered intragastrical. Blood glucose was measured before glucose load, at time 0, and after 15, 30, 60 and 90 min. Blood glucose was determined with a glucose meter (Accu Check, Roche, Switzerland) on blood samples collected from the tip of the tail vein. Plasma insulin concentration was determined using an ELISA kit (Mercodia) according to manufacturer’s instructions.

### RNA isolation and quantitative RT-PCR

Frozen tissue samples were dissociated in lysing/binding buffer of Multi-MACS cDNA kit (Miltenyi) with 0.5% antifoam using the gentleMACS™ Octo Dissociator (Miltenyi). RNA isolation was performed using the PureLink RNA Mini Kit (Invitrogen) and cDNA synthesis was performed using the superscript IV (Invitrogen). Real-time quantitative PCR was performed using SybrGreen (BioRad) and RT^2^ qPCR primer assays (Qiagen) or PrimePCR^TM^ (Biorad) primers. Ct values were normalised to the Ct mean of three housekeeping genes (*Gapdh*, *Hsp90* and *Hprt*, or *Hprt*, *B2m* and *Tbp*) in each sample.

### Glucose absorption test

A glucose solution (1.5g/kg) (CDM Lavoisier) containing 10% 2NBDG (Invitrogen) was administered intragastrically; 15min post administration, intestinal segments were harvested and weighted. Samples were dissociated with beads using a tissue lyzer in 200µl of PBS. After centrifugation, supernatants were collected and fluorescence was read with an excitation wavelength of 465nm and an emission of 540nm using a fluorescence plate reader (SLT, BMG Labtech). Autofluorescence was determined using samples from mice administered intragastrically with glucose only.

### Single cell RNA sequencing of intestinal epithelial cells

Small intestinal epithelial cells (IECs) were isolated from wild-type (WT) and ROD3 mice under defined metabolic and treatment conditions. In the first experiment, IECs were collected from two WT mice maintained on *ad libitum* feeding, two WT mice following 16 h of fasting, and two ROD3 mice following 16 h of fasting. All mice in this cohort received an intraperitoneal injection of diphtheria toxin (DT) one day prior to tissue collection. In a second single cell RNA sequencing experiment, all mice received an intraperitoneal injection of DT one day before tissue collection and were subjected to 16 h of fasting prior to the experiment. Small intestinal epithelial cells were isolated from two WT mice treated with PBS (fasted control), two WT mice treated with CO lipid/LPS (fasted with stimulation), two ROD3 mice treated with PBS (fasted control), and two ROD3 mice treated with CO lipid/LPS (fasted with stimulation). In addition, IECs were collected from two WT and two ROD3 mice maintained on *ad libitum* feeding as non-fasted controls. Single-cell RNA sequencing was performed on the isolated epithelial cells from each group.

The small intestine was dissected and cleaned of feces and fat. The epithelial layer was dissociated by incubating the tissue twice for 15 min in HBSS supplemented with 5 mM EDTA and 10 mM HEPES at 37°C. Cells were then incubated in a digestion mix containing HBSS with Ca^2+^/Mg^2+^, dispase (5 U/ml) (BD Biosciences), collagenase D (0.5 mg/ml) (Roche), DNaseA (0.5 mg/ml) (Sigma-Aldrich), and 3% FCS during 10 mins at 37°C and then filtered on 70µm filter to prepare a single cell suspension. Cells from the two samples of each group were pooled for the subsequent steps. Single-cell suspensions were stained with EPCAM-APC (clone: G8.8), CD45- FITC (clone: 104), and DAPI to exclude dead cells. Epithelial cells were sorted using a FACS Aria II Cell Sorter (BD Biosciences). Approximately 6,000 cells per genotype, before quality control, were captured using the 10X Genomics Chromium Next GEM 3’ Library & Gel Bead Kit V3 (10X Genomics), according to the manufacturer’s protocol. Libraries were sequenced on an Illumina NextSeq 2000 P3 (138 cycles). Raw sequencing data were processed using Cellranger-3.0.2 (10X Genomics) with the mm10 reference genome and default parameters.

### Data Analysis

UMI counts were analyzed using Seurat 5 and integrated using Harmony. Variable genes were identified, and data were scaled and SCT transformed using the default parameters. UMAP and PCA were performed based on 30 components and a perplexity of 30. Clustering was determined using a resolution of 0.2. Clusters with low *Epcam* expression were excluded, and the analysis was repeated. After quality control and filtering we obtained in total (after integration of all conditions) 5092 cells for control *ad libitum*, 4305 cells for control fasted condition, 5681 for the ROD3 *ad libitum,* 3059 cells for ROD3 fasted condition, 5635 cells for control lipid/LPS and 6452 cells for ROD3 lipid/LPS. Differential expression analysis was conducted using Seurat 5, identifying genes with a log fold change > 2.5 and a minimum cell expressing set at default (0.1).. Genes were functionally categorized using an AI-assisted LLM-based classification across intestinal regions.

### Crypt Isolation and organoids generation

Intestinal crypts were extracted as previously described (*72*), embedded in Cultrex Ultimatrix (R&D) diluted 2:1 with culture medium, and 4 drops of 30 μl (containing 500 crypts each) were seeded in 12-well plates, incubated for 15 min at 37°C, and overloaded with 800 μl of Culture medium. The culture medium was composed of Advanced DMEM/F-12 or DMEM low glucose, supplemented with 100 U/ml penicillin/streptomycin, 10 mM Hepes, 1x N2, 1x B27 (all from Gibco), 50 ng/ml EGF (R&D), 100 ng/ml Noggin (R&D) and 500 ng/ml R-spondin1 (R&D). After four days of culture, the organoids were collected in tubes, spun at 300g at 4°C, and lysed with Lysis buffer from PureLink RNA mini Kit (ThermoFisher) before being processed for RNA extraction according to the manufacture instructions.

### RNAscope histology of organoids

For RNAscope experiments, 300 crypts were embedded in 18 μl of diluted Matrix and seeded in μslide 18-well glass bottom (Ibidi) covered with 100 μl of culture medium. After 4 days of culture, 100 μl of 8% PFA (Electron Microscopy Sciences) was added to the wells and fixed o/n at 4°C before proceeding to RNA-fluorescence in situ hybridization (RNA-FISH) using the RNAscope Multiplex Fluorescent v2 kit (Advanced Cell Diagnostic), adapting the manufacturer’s recommendations as follows. Samples were delipidated with 4%SDS-200mM Boric Acid 2h at RT, permeabilized with 0.5% triton-X100/PBS, washed with wash buffer (0.2% Tween-20, 20 µg/ml heparin in PBS) and incubated 10 min with 3% H_2_O_2_ in PBS at RT before hybridization with the target probes Lgr5 and Slc40. Following the amplification steps, the signal development for the two probes was performed sequentially using TSA Vivid 570 and 650 kits (Tocris) (respectively for Lgr5 and Slc40 probes). Incubation with E-cadherin (ECCD-2) antibody was performed o/n at 4°C followed by incubation with goat anti-Rat AF488 and 1 μg/ml DAPI. Images were acquired with the confocal microscope BC43 (Andor) using an objective 60x/1.42, deconvoluted using ClearView, and rendered in Imaris. The single stack images were generated with FiJi.

### Measurement of Reactive Oxygen Species in Fresh Intestinal Epithelial Cells

Fresh small intestinal epithelial cells were isolated from mouse gut as described above. Immediately after isolation, cells were resuspended in assay buffer and counted using an automated cell counter with fluorescent viability dyes (Logos Biosystems). For intracellular ROS levels assessement, freshly isolated epithelial cells were plated in black 96-well plates at densities of 25,000, 50,000, and 100,000 cells per well in 100 µL of buffer. Cells were incubated with 5-10µM of the fluorescent probes 2’,7’- dichlorofluorescin diacetate (DCFDA) and dihydrorhodamine 123 (DHR123), and 2-5 µM MitoSOX Red, for 20-30 min at 37 °C in the dark. After incubation, cells were washed three times to remove excess dye and fluorescence was measured using a fluorescence plate reader (SLT, BMG Labtech). Fluorescence intensity was recorded as relative fluorescence units (RFU) and normalized to viable cell number.

### Statistical analysis

P-values were calculated with unpaired, two-tailed parametric, or non-parametric statistical tests as stated in the figure legends using GraphPad PRISM (version 10) software. Normality was tested using Shapiro-Wilk. Exact tests and P values are shown. P values > 0.05 were considered not significant (ns).

### Data availability

The data generated or analyzed during this study are included in the manuscript and its Supplementary Information files. Single cell RNA sequencing data are available in the GEO repository as GSE279711.

## ACKNOWLEDGEMENTS

We thank the Microenvironment & immunity lab, and the Stroma, Inflammation and Tissue Repair lab for discussion. We also thank the staff of the Central Animal Facility, Flow Cytometry platform, Histopathology core platform, Biomics platform and Mouse Genetics Engineering platform of Institut Pasteur for their help and commitment. This project was supported by grants from the Institut Pasteur, the Agence Nationale de la Recherche (ANR) No ANR-15-CE15-0014-01, the Fondation de la Recherche Médicale (FRM) DEQ20160334871 and the Biostime Institute for Nutrition and Care – BINC Geneva Foundation Funding Programs. S.C. was supported by a Marie Curie intra-European Fellowship, F. G. by a Walter Benjamin fellowship from the Deutsche Forschungsgemeinschaft. B.C.’s laboratory is supported by a Starting Grant from the European Research Council (ERC) under the European Union’s Horizon 2020 research and innovation program (grant agreement No. ERC- 2018-StG- 804135), ANR grants EMULBIONT (ANR-21-CE15-0042-01) and DREAM (ANR-20-PAMR-0002), and the national program “Microbiote” from INSERM.

## AUTHOR CONTRIBUTIONS

E.L. designed and performed experiments, interpreted the results and wrote the paper. F.G designed, performed experiments, analyzed the single-cell data and organoid experiments, interpreted the results and wrote the paper. S.C. designed, generated and characterized the transgenic ROD mice, and designed, performed, analyzed experiments and interpreted results. G.N. designed, performed and interpreted the results of organoid experiments and RNAscope imaging. J.M. performed, analyzed and interpreted depletion strategy experiments. S.D. performed experiments and transcript profiling. M.R. performed experiments and histology imaging analysis. B.C. supported the targeted microbiota analysis. F. L. generated the transgenic mouse models. F.D. designed and generated the transgenic animal models and performed experiments. G.E. directed the research, conceptualized and supervised the project, interpreted the results and wrote the paper.

## FIGURES and LEGENDS

**Fig. S1.**
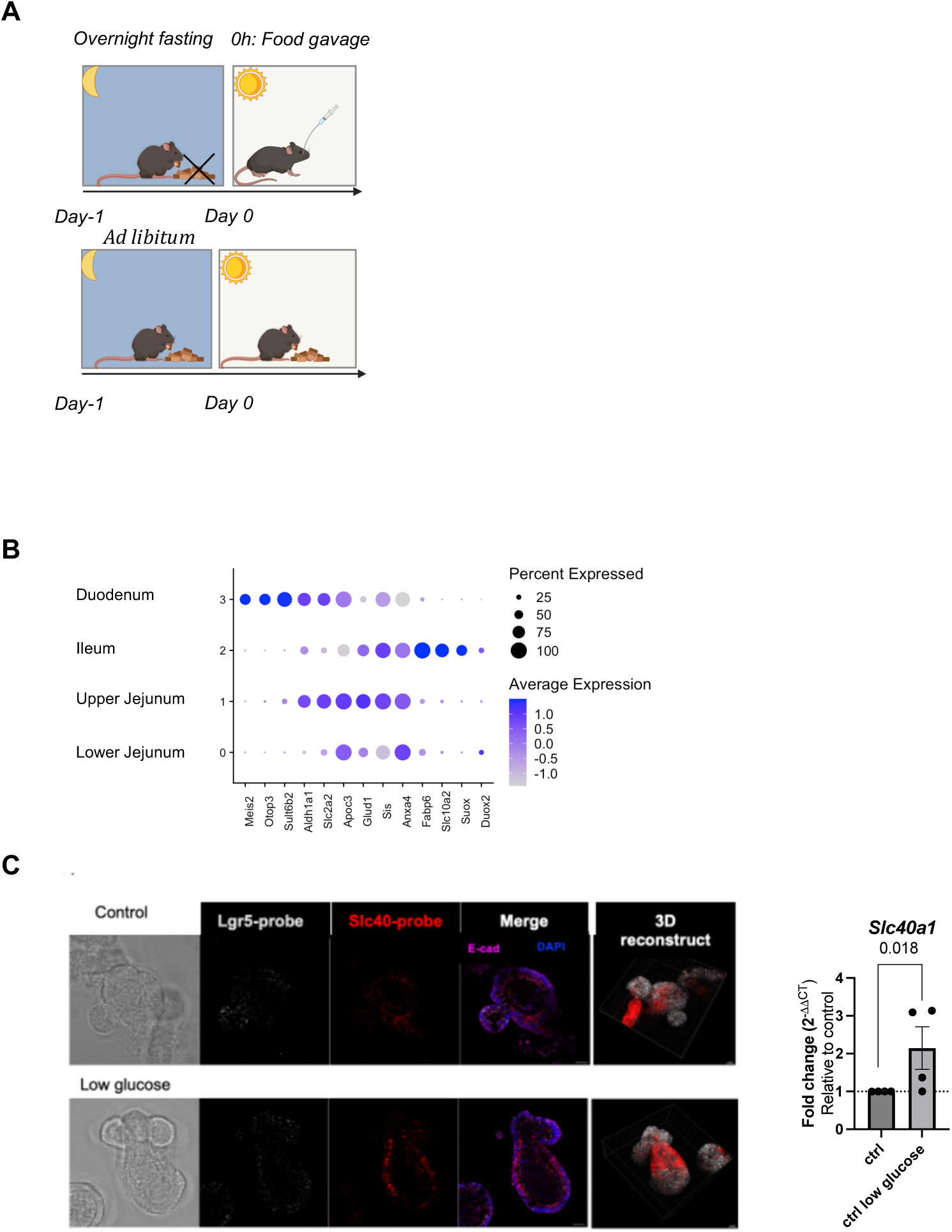
Fasting experimental setup and organoids. (**A**) Overnight fasted mice were gavage with different types of foods, or fed ad libitum, and analyzed at different time points after dietary challenge. (**B**) Dot plot of gene expression used to identify the different anatomical regions of the small intestine. (**C**) L*eft:* Representative RNAscope images of intestinal organoids probed for *Lgr5* (white), *Slc40a1* (red), E-cadherin (magenta) and stained for DAPI (blue) (scale bar 20µm). *Right*: Expression of *Slc40a1* (relative to control organoids) (mean ± SEM n = 4 organoid per group) in organoids grown in the indicated condition. Data in (B) are representative of one single cell RNA sequencing experiment. Data in (C) are representative of a pool of two independent experiments and representative of more than three independent experiments and symbol indicates individual mice. *P* values were determined by two-tailed nonparametric Mann– Whitney U test. Exact *P* values are indicated on graphs.

**Fig. S2.**
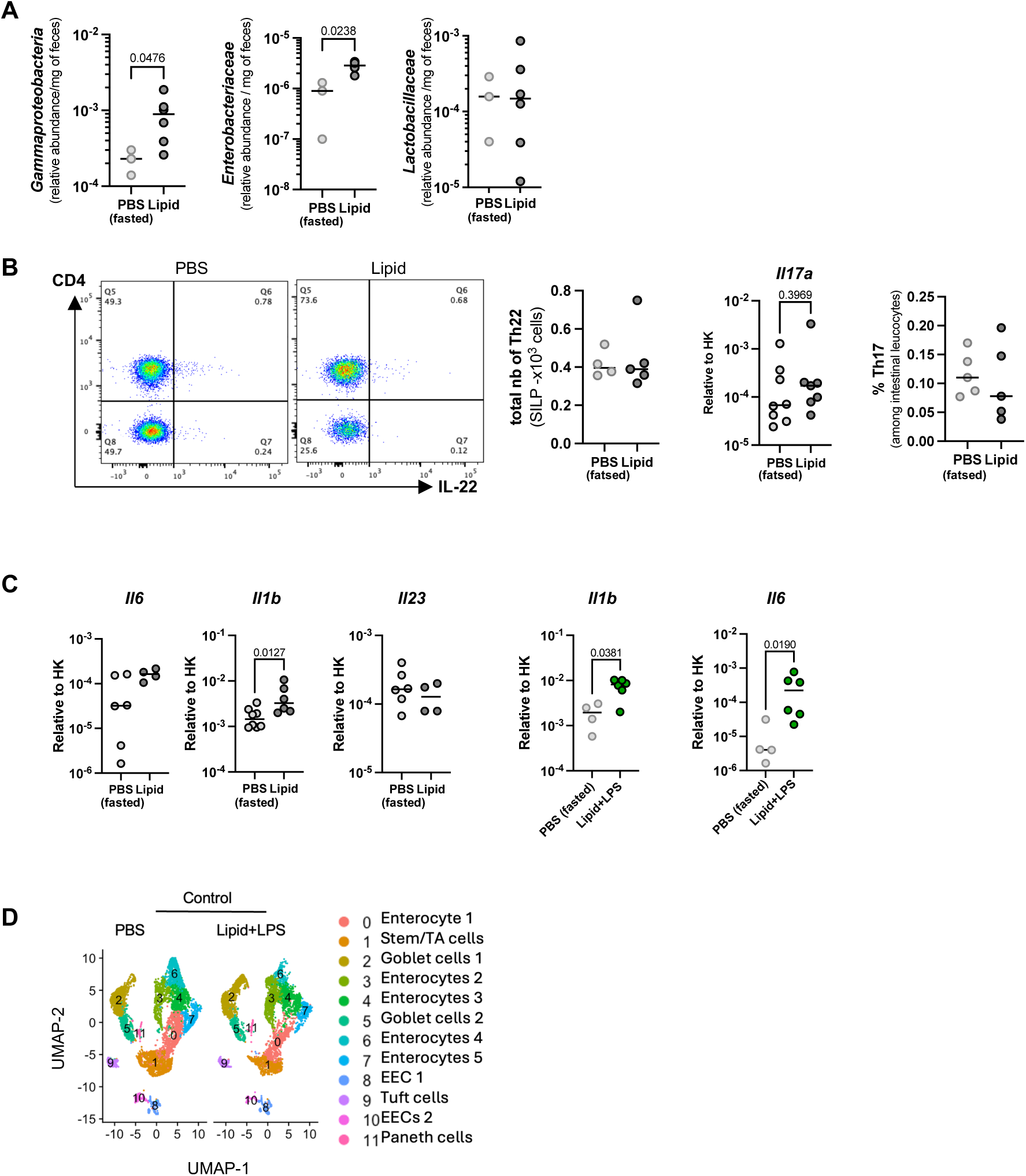
The effect of lipid-rich food intake on microbiota and intestinal immunity. (**A**) qRT-PCR (median, n = 3-6 per group) for γ*-Proteobacteriaceae*, *Enterobacteriaceae* (median, n =3-6 per group) and *Lactobacillaceae* (median, n = 3-6 per group) targeting 16S rRNA genes in feces of mice 90 min after CO lipid challenge. (**B**) *Left:* representative dot plots of small intestinal T cells gated on viable CD45^+^CD3^+^ cells and quantification (median, n = 4-5 per group) of absolute numbers of IL-22^+^ T cells, *middle*: relative expression of *Il17a* from ileal biopsies, *right:* percent of small intestinal Th17 cells gated on viable CD45^+^CD3^+^ of mice fed CO lipid or PBS (median, n = 5 per group) (**C**) Expression of *Il6*, *Il1b* and *Il23* measured by RTqPCR from ileal biopsies in control mice 90 min after dietary challenge with PBS (light grey), CO lipid (dark grey) or CO lipid/LPS (green). (**D**) UMAP projection of scRNA-seq data of enterocytes from mice fed PBS or CO lipid/PBS. Colors indicate unsupervised clustering determined by Seurat; clusters were annotated using canonical markers of enterocytes. Data in (A) to (C) are a pool of two or three independent experiments and representative of three or more independent experiments and symbol indicates individual mice. Data in (D) is from one single cell RNA sequencing experiments with a pool of 2 mice per group. *P* values were determined by two-tailed nonparametric Mann–Whitney U test or Kruskal–Wallis test. Exact *P* values are indicated on graphs.

**Fig. S3.**
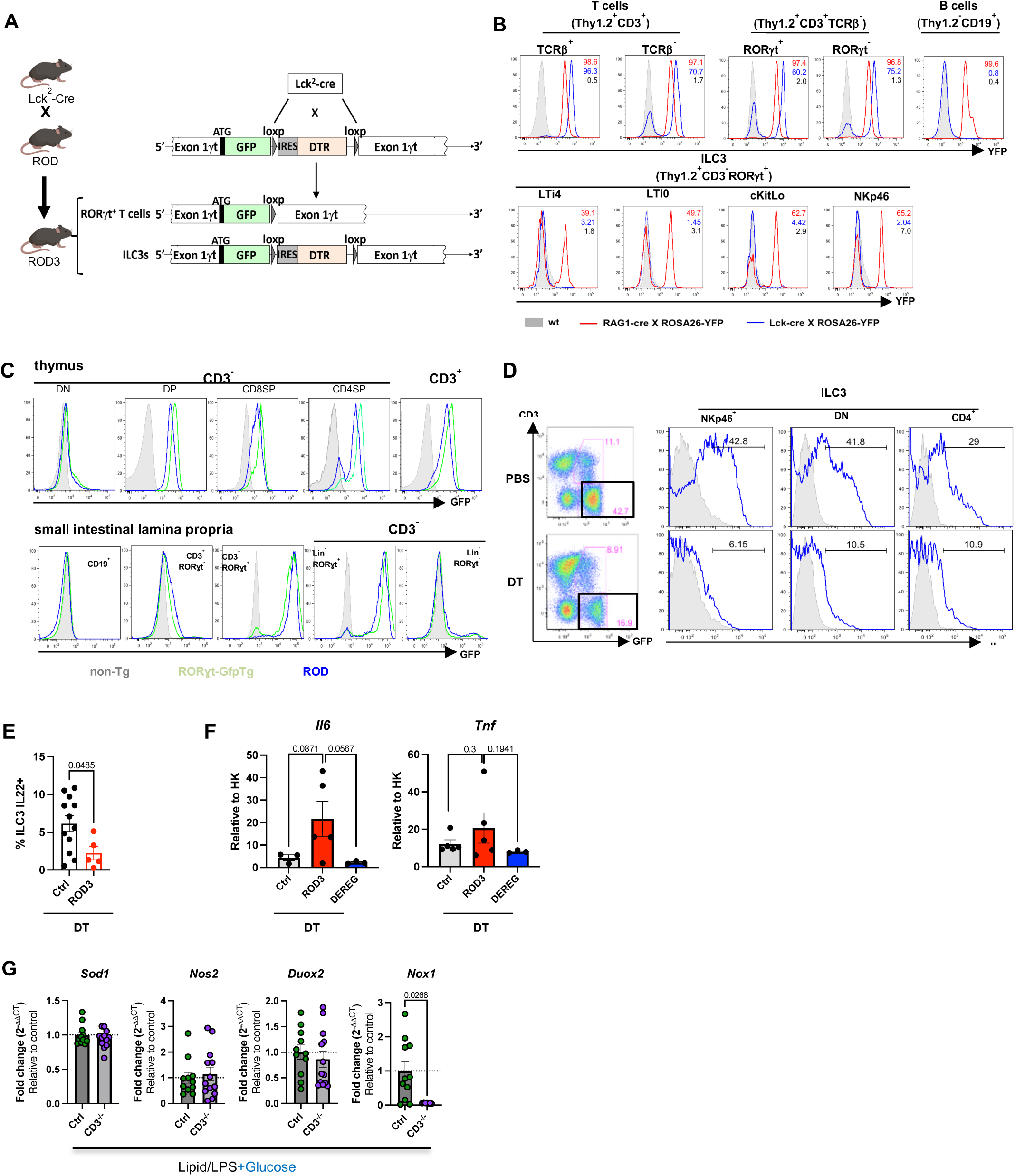
ILC3 depletion model – the ROD3 mouse. (**A**) Schematic of the gene construct in ROD mice, and mating strategy between the RORɣt-Depleter (ROD)-mice with Lck^2^-Cre mice to obtain the ILC3-depleter (ROD3) mice. (**B**) Histograms of flow cytometry analysis comparing YFP expression by small intestinal lamina propria cells isolated from Lck^2^-Cre (blue line) or RAG1-cre (red line) mice crossed to ROSA-YFP or from control (wt) mice (light grey). (**C**) Histograms of flow cytometry analysis comparing GFP expression by thymus (upper panel) and small intestinal lamina propria (lower panel) immune cells from RORɣt^GFP/+^ transgenic mice (green line), ROD^Tg/+^ transgenic mice (blue line) and control mice (light grey). (**D**) Representative CD3 versus GFP dot plot analyses of small intestinal lamina lymphoid cells (left), and representative histograms of ILC3s subtypes (right) for IL22 expression (blue line) isolated from the small intestinal lamina propria of ROD3 PBS-injected (upper panels) or diphtheria toxin (DT)-treated (lower panels) mice. Abbreviations: double negative (DN), double positive (DP), transgenic (Tg). (**E**) Quantification of ILC3-IL22^+^ cells in ad libitum control and ROD3 mice following overnight treatment with DT. (**F**) Expression (mean± SEM, n =3-6 per group) of *Il6* and *Tnf* in control, ROD3 and DEREG mice after overnight fasting and DT treatment, 90 min after feeding with CO lipid/LPS. (**G**) Relative expression (median, n = 7-11 per group) of *Sod1*, *Nos2*, *Duox2* and *Nox1* in control and *Cd3*^-/-^ deficient mice after overnight fasting and 105 min after feeding with CO lipid/LPS + glucose. Data in (E) to (G) are a pool of two to three experiments and representative of two or more independent experiments, and symbol indicates individual mice. *P* values were determined by two- tailed nonparametric Mann–Whitney U test or Kruskal–Wallis test. Exact *P* values are indicated on graphs.

**Fig. S4.**
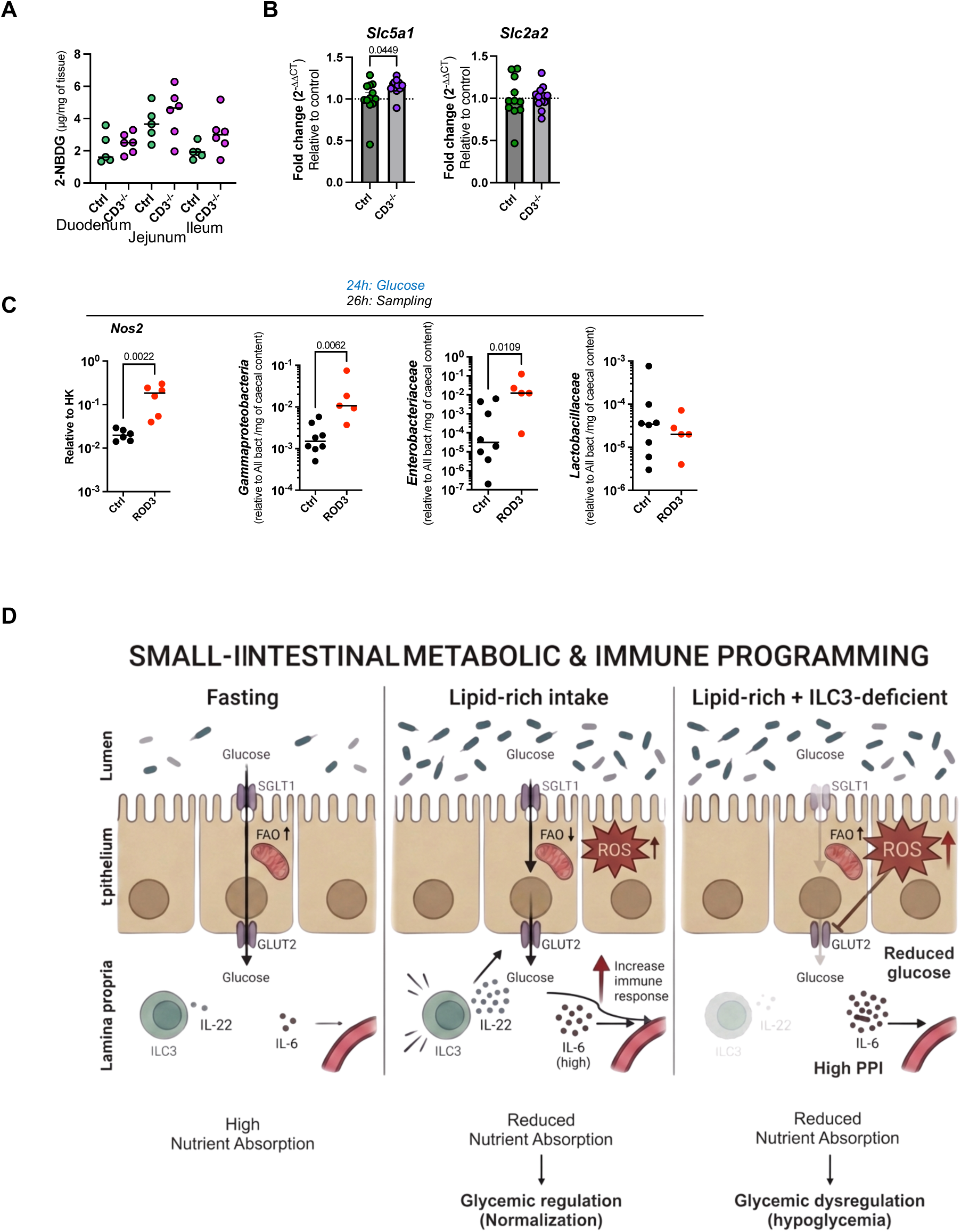
Glucose absorption in the absence of T cells, and graphical abstract. (**A**) Absorption (median, n = 5-6 per group) of 2-NBDG in the indicated intestinal compartment 15 min after oral gavage in control mice and *Cd3*^-/-^ mice previously fed CO lipid/LPS. (**B**) Relative expression (median, n = 10-11 per group) of *Slc5a1*and *Slc2a2* as determined by qRT-PCR in intestinal epithelial cells from control and *Cd3*^-/-^ mice 90 min after feeding with CO lipid/LPS. (**C**) *Left:* Relative expression of *Nos2* (median; n = 6–6 per group) determined by qRT-PCR in small intestinal epithelial cells from DT-treated control and ROD3 mice, and right: qRT-PCR (median, n = 5-8 per group) for γ*-Proteobacteriaceae*, *Enterobacteriaceae* and *Lactobacillaceae* targeting 16S rRNA genes in caecum of mice 2 hrs after glucose challenge performed 24h after CO lipid/LPS feeding. Data in (A) to (C) are a pool of two to three experiments and representative of two or more independent experiments, and symbol indicates individual mice. *P* values were determined by two- tailed nonparametric Mann–Whitney U. Exact *P* values are indicated on graphs. (**D**) Graphical abstract of the study.

